# Best practices for genome-wide RNA structure analysis: combination of mutational profiles and drop-off information

**DOI:** 10.1101/176883

**Authors:** Eva Maria Novoa, Jean-Denis Beaudoin, Antonio J Giraldez, John S Mattick, Manolis Kellis

## Abstract

Genome-wide RNA structure maps have recently become available through the coupling of *in vivo* chemical probing reagents with next-generation sequencing. Initial analyses relied on the identification of truncated reverse transcription reads to identify the chemically modified nucleotides, but recent studies have shown that mutational signatures can also be used. While these two methods have been employed interchangeably, here we show that they actually provide complementary information. Consequently, analyses using exclusively one of the two methodologies may disregard a significant portion of the structural information. We also show that the identity and sequence environment of the modified nucleotide greatly affect the odds of introducing a mismatch or causing reverse transcriptase drop-off. Finally, we identify specific mismatch signatures generated by dimethyl sulfate probing that can be exploited to remove false positives typically produced in RNA structurome analyses, and how these signatures vary depending on the reverse transcription enzyme used.

## INTRODUCTION

In the last decades, it has become clear that RNAs are not simply intermediaries between DNA and protein, but are in fact functional molecules capable of regulating central cellular and developmental processes, such as genome organization and gene expression, and comprise the bulk of human genomic programming ^1–4^. Because RNA is a single-stranded molecule, it tends to fold back on itself, forming stable secondary and tertiary structures by internal base pairing and other interactions. RNA structure plays an essential role in determining the function and dynamics of these molecules, and varies depending on environmental conditions ^5,6^. Thus, accurate genome-wide RNA structural maps can allow for better understanding of the complexity, function, and regulation of the transcriptome ^7,8^.

Dimethyl sulphate (DMS) and 2’-hydroxyl acylation and primer extension (SHAPE) reagents have traditionally been used to obtain experimental measurements of RNA structure, providing information on base-pairing and tertiary interactions of the RNA molecules ^9–11^. These chemicals show selective reactivity toward unpaired RNA bases. Until recently, limitations of probing reagents as well as in sequencing and informatics restricted structural profiling analyses to a few *in vitro* folded RNAs. However, the coupling of DMS and SHAPE chemical labeling with next-generation sequencing ^12–18^ now allows for creation of genome-wide RNA structure maps that provide substantial information regarding the dynamics of RNA structures in a variety of cellular contexts ^12,15^.

Initial attempts to analyze genome-wide RNA structure data relied on the truncation of reverse transcription upon reaching a nucleotide that has been chemically modified by probing reagents ^13,15,16,19,20^. However, this method suffers from several limitations and caveats, such as the presence of naturally occurring modified nucleotides, the addition of untemplated nucleotides by the reverse transcriptase (RT) ^21^, or the presence of stable structures leading to truncated reverse transcription, hindering correct annotation of the modified base. At least 8% of the annotated positions constitute false positives, as 8% of annotated read ends of DMS-Seq analyses are typically mapped onto G and T bases ^15^, while only A and C bases should be identified by this technique. Moreover, RNA and DNA ligases commonly used for adapter ligation have distinct sequence preferences for the ends being ligated, biasing the representation of particular read ends in the resulting libraries ^22^.

To overcome these limitations, several groups used the increased mismatch rates that occur at DMS- and SHAPE-modified nucleotides to chart RNA structure ^23^, known as mutational profiling (MaP), giving rise to SHAPE-MaP ^24^ and DMS-MaPSeq ^25^. Unfortunately, mutations occur at low frequency, and therefore can only be identified in transcripts with very high read coverage. Consequently, these methods are generally limited to the analysis of individual RNA transcripts ^23,24,26,27^, smaller genomes such as the HIV-1 RNA genome ^23^, or to highly expressed genes in more complex genomes ^25^.

Here we use both methodologies to analyze transcriptome-wide DMS-seq datasets of *in vivo* probed 64-cell stage zebrafish embryos (see Methods) —including well-characterized RNA structures as spike-ins— and compare the set of DMS-modified positions using either the reverse transcription truncation signals or MaP. Previous studies employed either reverse transcription truncation signals 15or MaP ^25^, with the underlying assumption that results largely overlap. Contrary to these expectations, we find a low correlation between the two methodologies, despite correct identification of unpaired nucleotides by each methodology. Furthermore, we observe that the probability of driving a mismatch or a RT-stop signal upon reverse transcription is dependent on the identity of the modified nucleotide, the surrounding sequence context, and the nature of the reverse transcriptase enzyme employed in the library preparation. Considering our findings, we suggest that RNA structural analyses based on DMS or SHAPE probing should combine both RT truncation signals and MaP to best capture structural information, and show that the accuracy of RNA structure predictions can be increased when both types of information are used.

## RESULTS

### Mutational profiling and RT drop-off analyses capture complementary information

DMS has traditionally been used to probe RNA structure *in vivo*. It reacts with the *N1* of adenosines, *N3* of cytosines and *N7* of guanosines in single stranded (ss) RNA, resulting in chemically modified nucleosides 1-methyladenosine (m^1^A), 3-methylcytosine (m^3^C) and 7-methylguanosine (m^7^G), respectively. Unlike m^7^G, m^3^C and m^1^A methylations affect the Watson-Crick base-pairing (Figure 1A), and can therefore cause reverse transcription (RT) drop-off ^28^ as well as increased error rates (‘mismatch’ patterns) upon reverse transcription ^29,30^ (Figure 1B).

**Figure 1.**
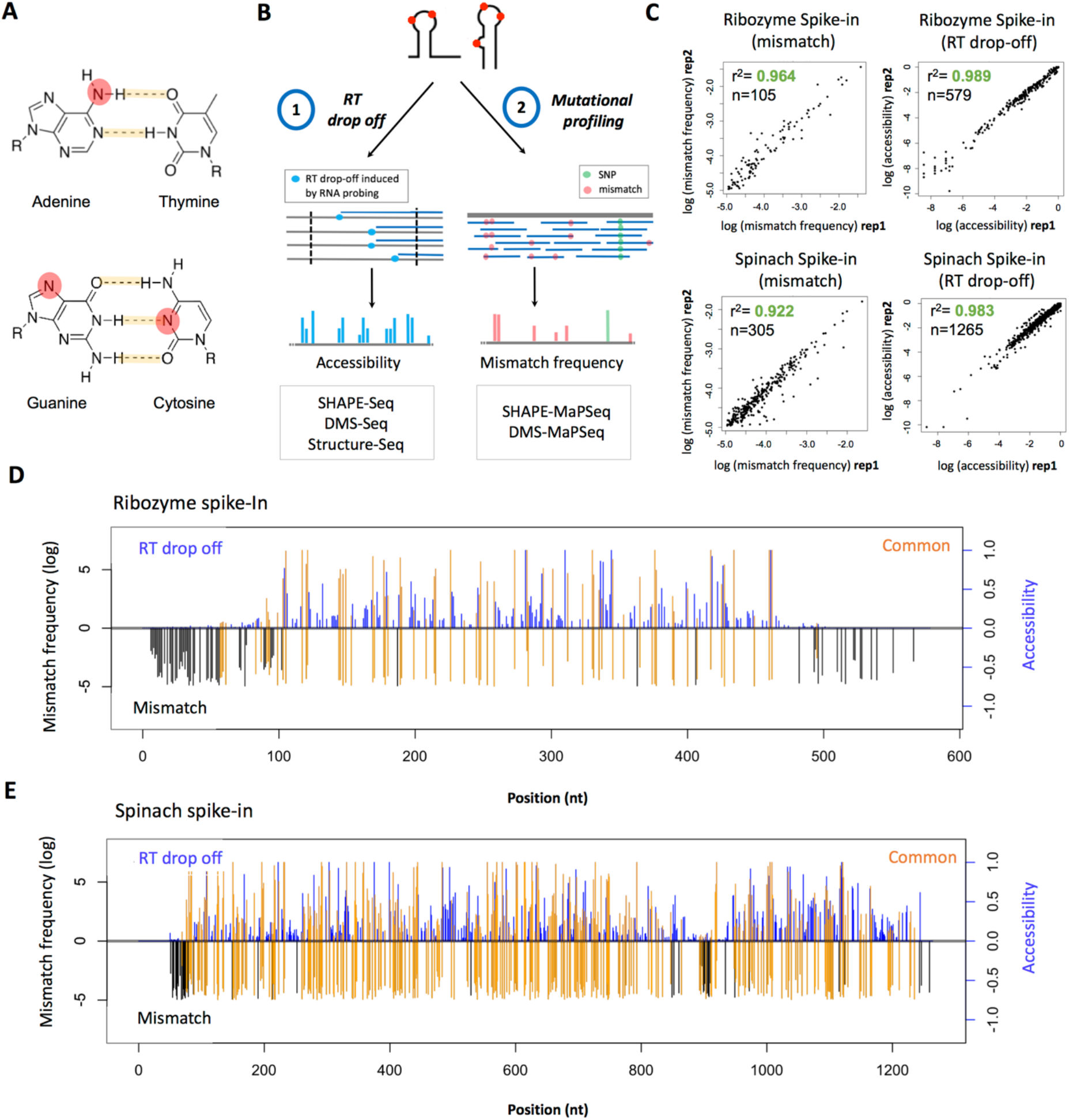
Mutational profiling and RT drop-off datasets contain non-overlapping information. (**A**) DMS methylates nitrogen atoms (red), some of which are involved in Watson-Crick base pairing (m^1^A and m^3^C), causing RT drop-off and increased mutational rates during reverse transcription. (**B**) Overview of the two major methodologies relying either on RT drop-off or mutational profiling to quantify DMS signal. (**C**) Replicability of mutational profiles and RT drop-off accessibilities of two RNA spike-ins. (**D and E**) Correlation between mutational profiles (“mismatch”, black) and RT stops (“accessibility”, blue) in the *Tetrahymena* ribozyme (D) and the DsRed mRNA containing a spinach tRNA cassette in its 3’UTR (E) spike-ins. Overlapping positions (“common”) detected by both methodologies are depicted in orange.

To compare the set of DMS-modified positions identified using reverse transcription truncation methodology (RT drop-off) or mutational profiling methodology (MaP) (Figure 1B), we first examined the RT-stop signals and mutational profiles of two spike-ins with known RNA structure —the *Tetrahymena* ribozyme (Rz) ^31^ and the tRNA spinach cassette (Spi) ^32^. Read coverage for these two molecules was extremely high (~50,000X), facilitating the identification of low frequency mismatch positions observed upon DMS probing with high confidence. We found that both methodologies quantified the level of per-nucleotide DMS modification with high replicability in both spike-ins (Pearson’s r=0.92-0.99) (Figure 1C). However, when we compared the individual positions identified by each methodology, we found only a partial overlap between the two methods (Figures 1D and 1E), suggesting that mutational profiling and RT truncation methods are capturing non-overlapping sets of DMS-modified nucleotides.

Direct comparison of the identified positions shows that mutational profiling methods capture DMS-modified positions at the 5’ and 3’ end of the molecule (Figures 1D and 1E), whereas these positions are not identified by RT drop-off methodologies due to an intrinsic limitation of the library preparation, which includes a second size selection step. When comparing the overlap of positions identified by each methodology, we find that 40% of positions identified using mutational profiling are not identified by RT-stop methodologies in transcriptome-wide analyses (Figure 2A). Similarly, in the spike-ins, 11-26% of mismatched positions are not identified by RT methodologies (Figure 2B).

**Figure 2.**
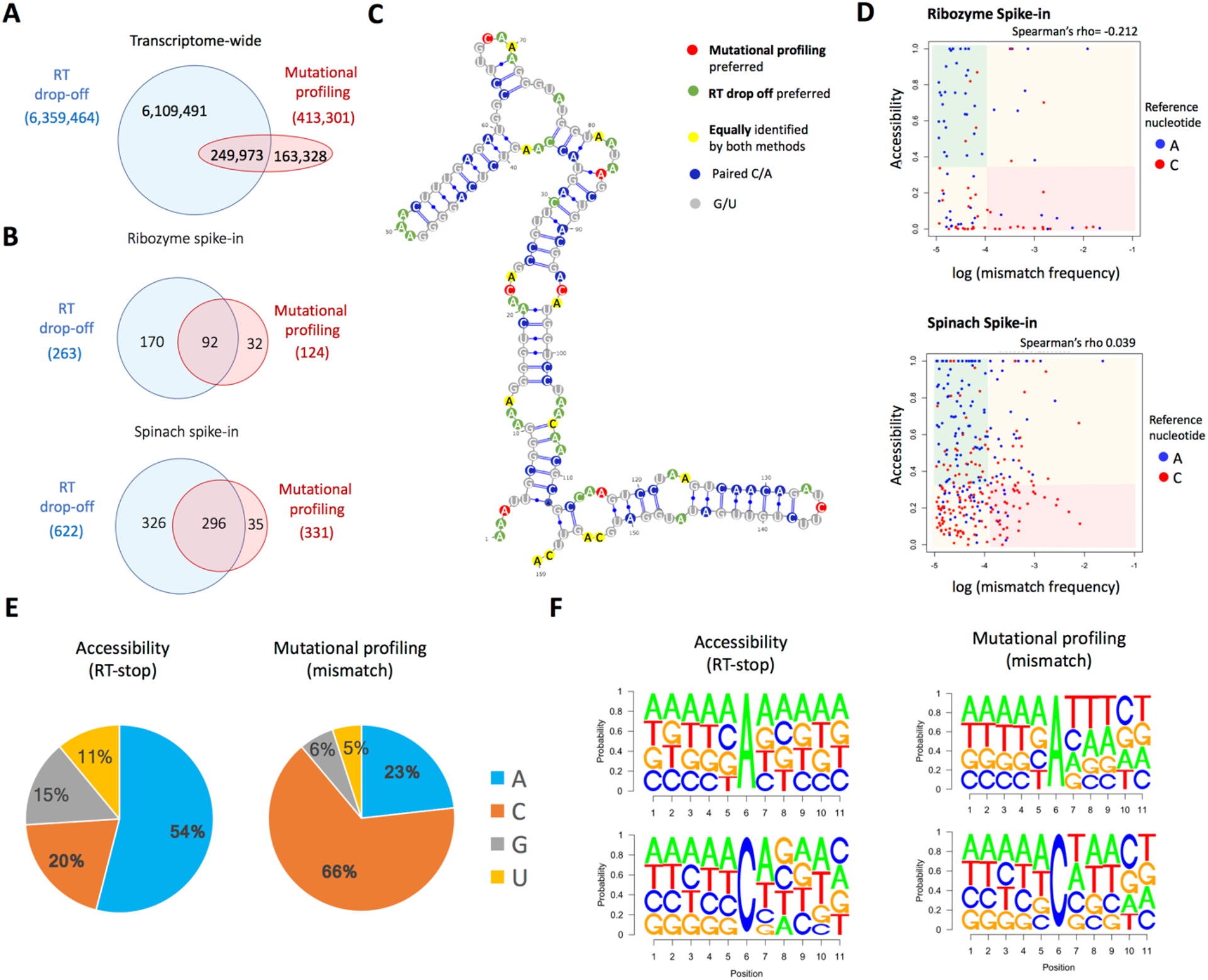
Both detection methods provide reliable structural information and are influenced by the nature of the DMS-modified nucleotide. (**A and B**) Overlap of DMS-modified positions identified using mutational profiling or RT truncation methodologies in the zebrafish transcriptome (A), and the *Tetrahymena* ribozyme (B, top) and DsRed mRNA with a spinach tRNA cassette (B, bottom) spike-ins. (**C**) Mapping of the mismatch frequencies (MaP) and the accessibilities (RT-stop) onto the experimentally determined RNA secondary structure of the *Tetrahymena* ribozyme P4-5-6 domain. Nucleotide positions preferentially identified by MaP are colored in red, those preferentially identified by RT stops are green, those equally identified by both approaches are yellow, and paired nucleotides are colored in blue. Gs and Ts, which are not detectable using DMS probing, are colored in gray. See also Figure S1. (**D**) Correlation between the accessibility and the mismatch frequency of the DMS-modified positions in the two spike-ins. Each nucleotide is colored according to its reference nucleotide. The area of the plot is divided into 3 regions: *i*) mismatch preference (high mismatch frequency, low accessibility), shaded in red; *ii*) RT-stop preference (high accessibility, low mismatch frequency), shaded in green; and *iii*) no preference, shaded in orange. (**E**) Proportion of reference nucleotides in transcriptome-wide DMS-seq experiments from 64-cell stage zebrafish embryos, using the RT-stop (left) or the mutational profiling (right) methodologies. (**F**) Comparison of the sequence context of mismatched positions and RT-stop positions in DMS-probed 64-cell stage zebrafish embryos.

The number of positions identified by the RT-stop signal methodology is ~2-fold higher than the mutational profiling methodology in the spike-ins (Figure 2B), and up to 15-fold higher when analyzing transcriptome-wide datasets (Figure 2A). This difference is partly due to the second size-selection step in the library preparation, which enriches the sample in RT truncated reads, as well as the coverage requirement of 50 reads/nucleotide to identify mismatched positions (see Methods), which limits the set of predicted mismatches to highly expressed genes.

To ascertain whether both methodologies correctly identified unpaired positions, we superimposed the predicted mismatch and RT-stop positions onto the known secondary structure of the *Tetrahymena* ribozyme ^31^, finding that both methods correctly identify unpaired positions (Figure 2C and **S1**). Intriguingly, we find that some modified positions are preferentially identified using the RT truncation methodology (green), whilst others are preferentially identified using mutational profiling (red) (Figure 2C).

### Choice of mismatch or RT stop is dependent on the nature of the DMS-modified nucleotide

We hypothesized that the nature of the modified nucleotides -m^1^A or m^3^C-might be partly dictating whether the reverse transcriptase incorporates a mutation or drops off. To test this, we compared the signal from both methodologies at individual As and Cs, in the spike-ins (Figure 2D) and transcriptome-wide (Figure 2E). We find that positions predominantly identified by the mutational profiling methodology were largely Cs, whereas those preferentially identified by the RT-stop method corresponded to As. Moreover, we observed low correlation between the two signals (Figure 2D), in agreement with previous works^27^.

Previous genome-wide studies employing RT truncation methodologies to retrieve DMS-modified positions had reported that 68% of the modified positions had an adenosine as the underlying nucleotide, while only 24% of the positions had a cytosine ^15^. Consequently, DMS has been generally thought to preferentially modify adenosines compared to cytosines ^1^. We find that RT-stops tend to occur more frequently at adenosines (54%), in agreement with previous reports ^15^. However, we observe that 66% of the mutational profiling signal arises from cytosines (Figure 2E), challenging the view that DMS preferentially modifies adenosines. Our results suggest that DMS does not favorably react with one nucleotide or another, but instead, that the method of analysis determines which DMS-modified positions are likely to be detected (Figure 2E).

We then investigated whether the sequence context was affecting the outcome of the reverse transcription, and if it could be responsible for the discrepancy observed between the two methodologies. We find that sequence contexts using either methodology appear to be slightly enriched in AT nucleotides (Figure 2F), which could be due to biases introduced during library preparation ^33^. When comparing the two methodologies, we observe no differences in the sequence context in the 5’ vicinity of the modified positions. In contrast, we do observe modest differences in the 3’ sequence context (Figure 2F). At first sight, our results suggest the sequence context may be playing a role in determining the fate of the reverse transcription (i.e., mismatch or RT-stop). However, library preparation biases, such as ligation bias, will also affect the sequence context identified using RT-stop methodologies, but not the sequence context identified using mutational profiling. Consequently, we cannot exclude the fact that the observed sequence context differences may partially arise from library preparation biases unequally affecting the two methodologies.

### Coupling RNA-Seq to DMS-Seq increases mutational profiling signal-to-noise ratio

To understand the discrepancy between MaP and RT-stop methodologies, we investigated whether false positives could be in fact biasing our results. For this aim, we analyzed the mismatch frequencies across replicates using 64-cell zebrafish embryo DMS-Seq datasets (Figure 3A, top panel). For comparison, we analyzed mismatch frequencies from 64-cell zebrafish embryo RNA-seq datasets (Figure 3B, top panel). As could be expected, we find that most mismatches identified in RNA-Seq datasets have mismatch frequencies ~1, suggesting these positions correspond to single nucleotide polymorphisms (SNPs). In contrast, mismatches identified in DMS-Seq datasets display a bimodal distribution, where some have mismatch frequencies ~1 -corresponding to SNPs-, whereas a second population displays very low mismatch frequencies. By comparing the patterns of RNA-Seq and DMS-Seq datasets, it is clear that low frequency mismatches observed in DMS-Seq datasets are the ones of interest, i.e., appearing upon DMS treatment. Therefore, we discarded data from positions with mismatch frequency greater than 0.25 in both the DMS-Seq and RNA-Seq datasets (Figures 3A and 3B, bottom panels). Surprisingly, there was still a significant number of mismatched positions present in RNA-Seq datasets. Most of these mismatches were consistent across RNA-Seq replicates, and were also found in DMS-Seq datasets (Figure 3C), suggesting that these may represent naturally occurring RNA modifications or low frequency SNP alleles in our population of embryos, which can act as confounders and should therefore be discarded.

**Figure 3.**
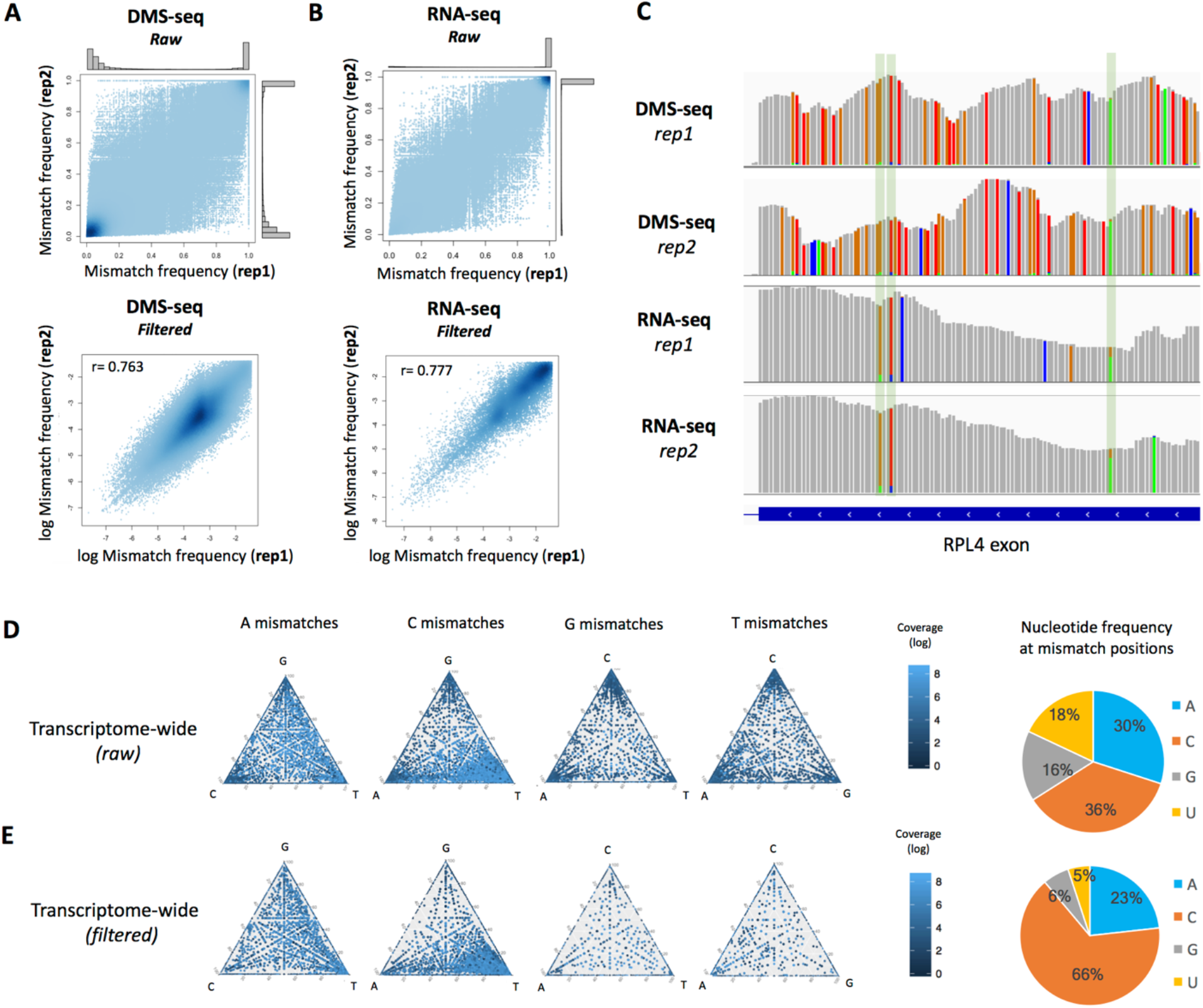
Filtering the noise from DMS-induced mismatch signal in a vertebrate transcriptome. (**A and B**) Replicability of mismatch frequencies for both DMS-Seq (A) and RNA-Seq (B) datasets, using raw mismatches (no filtering, top panels) and filtered mismatches (bottom panels, see Methods). (**C**) Mismatch positions (frequency > 0.01) along exon 9 of the RPL4 gene, in DMS-Seq and RNA-Seq datasets. Mismatches identified in each individual replicate are shown. Positions with mismatch frequency greater than 1% in both RNA-seq and DMS-Seq are highlighted, using IGV default coloring schemes. Mismatched positions that are consistent across DMS-Seq and RNA-Seq replicates are highlighted with green rectangles. (**D and E**) Ternary plots highlighting the mismatch signatures found in the DMS-modified transcriptome. For each reference nucleotide, the relative proportion of the other three nucleotides at mismatched positions is shown. The top (D) and bottom (E) rows depict transcriptome-wide DMS-Seq mismatches found in zebrafish 64-cell stage embryos, pre- and post-filtering. See also Figure S3 and Methods. The right pie charts show frequencies of reference nucleotides at mismatch positions.

### DMS probing generates distinct mismatch signatures for m^3^C and m^1^A

Previous works have suggested that some RNA modifications cause non-random misincorporation of nucleotides upon reverse transcription, known as ‘mutational signatures’ ^29,30,34^. Therefore, we hypothesized that m^1^A and m^3^C mismatches caused by DMS probing should generate identifiable mutational signatures. To identify the substitution patterns that occur upon DMS treatment in 64-cell zebrafish embryos, we compared the relative frequencies of misincorporated nucleotides at mismatch positions (Figure 3D), subdivided based on their reference nucleotide. From the ternary plot representations, we observed that both A and C mismatch positions did not randomly incorporate nucleotides (which would lead to scattered accumulation of points near the triangle’s center), but rather showed a biased “signature”, introducing mismatched nucleotides at specific frequencies (**Figure S2**). More specifically, m^1^A nucleotides preferentially misincorporated G and T bases, while m^3^C nucleotides favors misincorporation of T bases (Figures 3D and **S2**).

Upon examining all the mismatched positions, with no filter, we found that the majority (66%) of mismatches were found at A and C bases (Figure 3D). However, we also identified mismatches at G and U bases, which are not generated by DMS modification and should therefore not be observed. We therefore discarded: i) positions with mismatch frequencies higher than 0.25, ii) positions that are present and reproducible in RNA-Seq datasets, and iii) mismatches with insufficient coverage or with very low frequency of mismatches, which are likely PCR/sequencing artifacts (see Methods and **Figure S3**). Applying these filters removed most of the mismatches observed at G and T bases (Figure 3E), supporting the validity of pairing RNA-Seq with DMS-Seq for mutational profile analysis of RNA structure datasets. The inclusion of this filtering step led to 89% of the underlying nucleotides being A and C bases (Figure 3E). This updated mutational signature meets expectations of a profile generated by a single RNA modification (m^1^A and m^3^C, respectively), and supports the filtering steps as means to increase the signal-to-noise ratio, despite loss of some true positives (**Figure S3**), allowing for correct identification of DMS-specific mutational profiles (Figure 3E).

### Different reverse transcriptase enzymes show distinct misincorporation rates at m^3^C and m^1^A positions

The retroviral reverse transcriptase SuperScript-III enzyme (SS3) is the most commonly used RT enzyme in next-generation sequencing library preparations, including RNA-Seq and DMS-Seq. However, in the presence of modified RNA nucleotides that affect Watson-Crick base pairing, retroviral reverse transcriptases lead to a large proportion of RT drop-off compared to mismatched nucleotide incorporation ^25,35^. In contrast, thermostable group II intron reverse transcriptase (TGIRT) enzymes have higher processivity, fidelity, and thermostability than retroviral RTs ^36^, leading to lower RT drop-off rates ^35^ and increased mismatch frequencies ^25,27^.

Previous studies pioneering DMS-MaPSeq employed TGIRT to increase the proportion of mismatches in their datasets, and identified 51% of the DMS-modified positions to be adenosines ^25^. This finding is in contrast with our results using SS3, which identify cytosines at 66% of DMS-modified positions, and only 23% at adenosines (Figure 2E). We hypothesized that the choice of reverse transcriptase enzyme may not only increase the number of mismatches, but also unequally increase the mismatch frequency at m^1^A-modified positions compared to m^3^C. To test this, we used both SS3 and TGIRT for reverse transcription on DMS-modified 64-cell zebrafish embryos, employing a Structure-Seq protocol (see Methods). We find that the choice of enzyme used for reverse transcription has a significant effect on the proportion of reference nucleotides in DMS-modified samples (Figure 4A).

**Figure 4.**
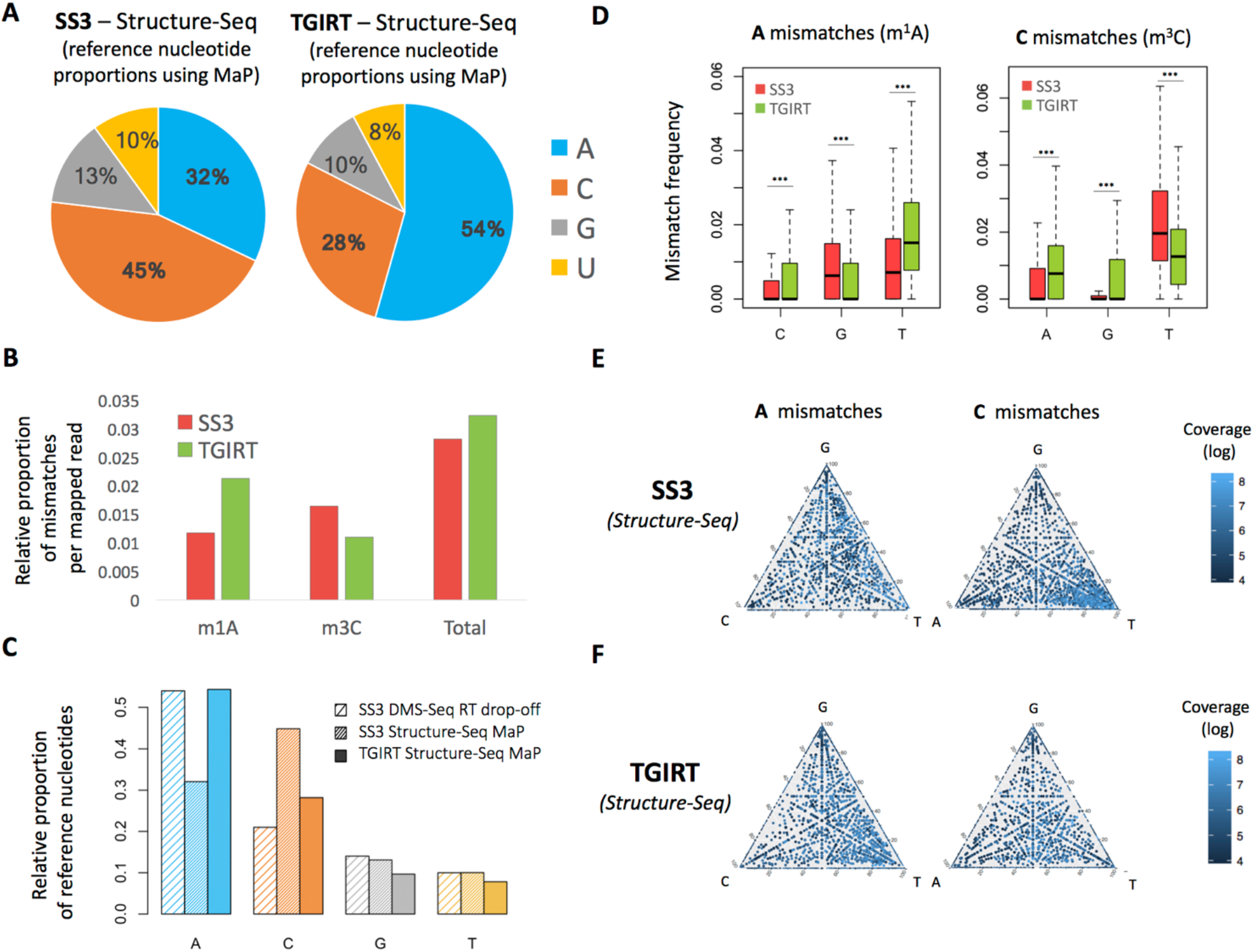
Mutational profiling signatures differ between reverse transcriptases. (**A**) Proportion of reference nucleotides with mismatches in transcriptome-wide Structure-seq experiments using either the SuperScript-III (SS3, left) or TGIRT (right) reverse transcriptases, with DMS-modified positions identified using mutational profiling. (**B**) Proportion of mismatched positions identified with the SS3 and the TGIRT reverse transcriptases. (**C**) Comparison of the relative proportion of reference nucleotides, when using different reverse transcriptases and analytical methodologies. (**D**) Proportion of misincorporated nucleotides at mismatched positions, using either the SS3 or TGIRT reverse transcriptase, at m^1^A (left) and m^3^C positions (right). Boxplot outliers have been removed for enhanced visualization. *P-*values were computed using Wilcoxon signed rank test (**E and F**) Ternary plots showing the transcriptome-wide mismatch signatures induced by m^1^A (left) and m^3^C (right), when using either the SS3 (E) or the TGIRT (F) enzymes in Structure-Seq datasets.

In agreement with this observation, we find that the increased processivity of TGIRT is not equal across DMS-modified positions; m^1^A positions exhibit higher processivity than m^3^C positions (Figure 4B). Consequently, the mismatch signal from the TGIRT experiment was enriched at adenosines, whereas the SS3 experiment displayed a stronger signal at cytosines (Figure 4A). Interestingly, we find the relative proportions of reference nucleotides of SS3 samples analyzed using the RT-stop methodology are similar to those reverse transcribed using TGIRT and analyzed using MaP (Figure 4C). This coincidental similar proportion of relative reference nucleotides may explain why prior works had not further characterized the non-overlapping nature of the two methodologies.

### The choice of reverse transcriptase enzyme impacts the mutational signatures

It has been suggested that different RNA modification may produce different mismatch ‘signatures’, where the identity and relative proportion of the misincorporated nucleotides may provide a clue to the nature of the underlying RNA modification ^29,30^. Indeed, previous works have employed machine learning algorithms trained with known tRNA modifications to predict the nature of each RNA modification based on its mismatch pattern ^29^.

As the relative misincorporation rate is affected by the RT enzyme used (Figures 4A-C), we wondered whether the identity of the misincorporated nucleotides, i.e., the ‘mutational signature’, was also different when using different RT enzymes. In this regard, we find that the mutational signatures are similar in datasets that used SS3 (Structure-Seq and DMS-Seq) (Figures 4D and 4E, compared to Figure S2D and Figure 3E, **respectively**). In contrast, the mismatch signature drastically changes when using the TGIRT enzyme. Specifically, m^1^A favors a substitution for a thymine while m^3^C doesn’t exhibit the clear preferential mutational signature seen with SS3 (Figure 4D and 4F).

Altogether, we find that the RT enzyme used for library preparation alters both the frequency and identity of the nucleotides that are misincorporated. Hence, we propose that mismatch signatures are specific to each RNA modification-reverse transcriptase combination, and not an intrinsic property of RNA modifications.

### Mutational signatures can enhance the accuracy of mutational profiling predictions

We have shown that DMS-dependent m^3^C and m^1^A RNA modifications have specific mutational signatures (Figures 3E and **S2**), and that these signatures are dependent on the RT enzyme employed in the library preparation (Figure 4D-F). However, certain RNA molecules, such as ribosomal RNAs, already contain a significant proportion of modified RNA nucleosides, which can confound the analysis. We therefore wondered whether the identified mismatch signatures could be used to discriminate DMS-probed positions (‘true positives’, TP) from those that are unrelated to DMS probing, such as naturally occurring RNA modifications (‘false positives’, FP). We used previously published DMS-probed *S. cerevisiae* datasets ^15,25^, which include yeast rRNAs with experimentally determined structures ^37^, to compare the mismatch signatures of “paired” (double stranded) versus “unpaired” (single stranded) residues (Figure 5A). We find that the mutational signatures alone (represented as T/G ratio for m^1^A mismatches and T/A ratio for m^3^C mismatches) do not separate the two populations (paired and unpaired) into different clusters effectively (Figure 5B, lateral density plots) (coefficients of overlap of the two distributions are 0.626 (m^1^A) and 0.769 (m^3^C)). Similarly, the mismatch frequencies alone do not separate them either (Figure 5B, top density plots) (coefficients of overlap of the two distributions are 0.726 (m^1^A) and 0.664 (m^3^C)). In contrast, we find that the combination of mutational signatures and mismatch frequencies effectively separates the paired and unpaired nucleotide populations (Figure 5B) (coefficients of overlap of 0.473 (m^1^A) and 0.306 (m^3^C)), suggesting that a combination of both types of information can increase the accuracy of RNA pairing status predictions.

**Figure 5.**
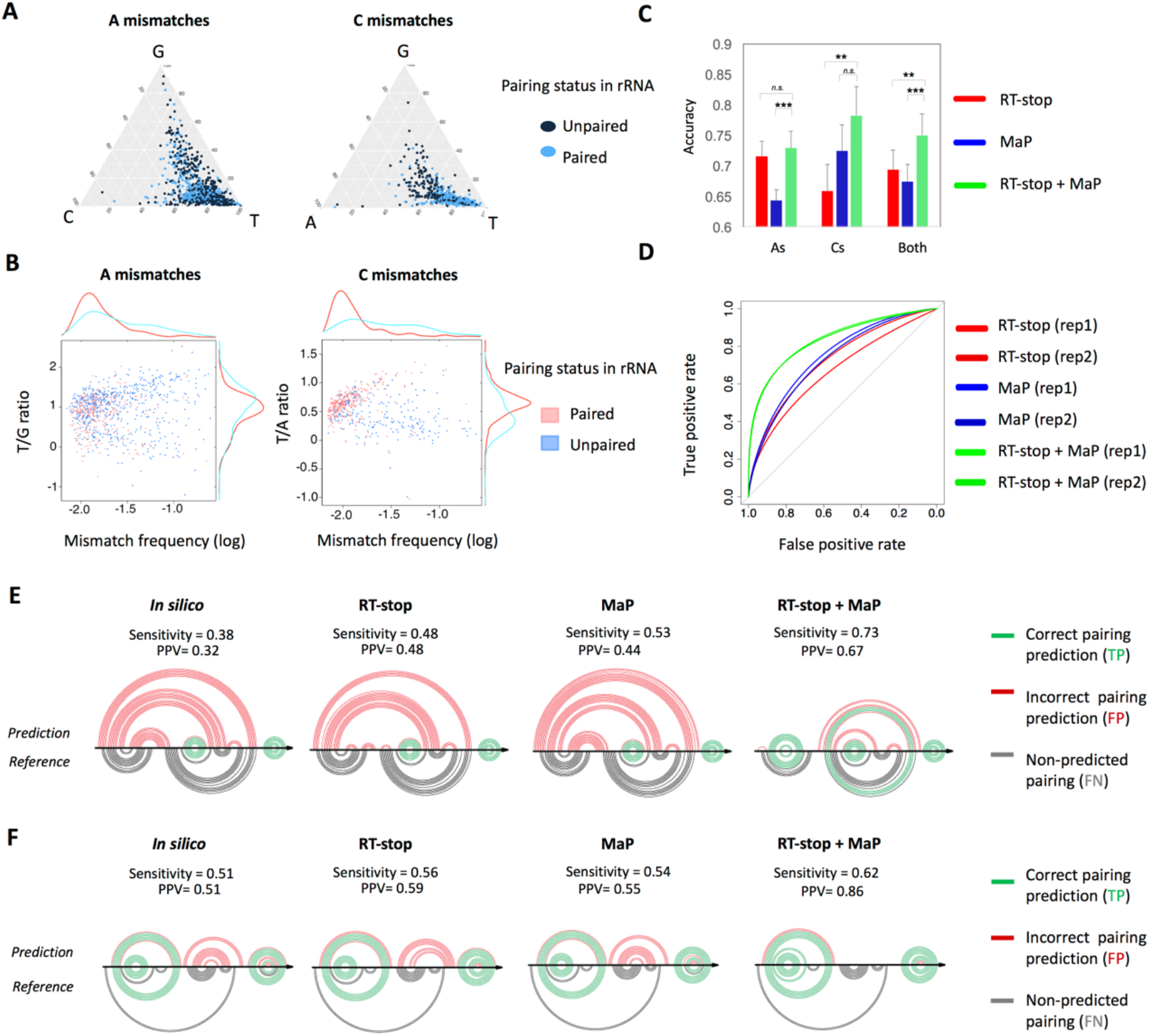
Combination of methodologies improves RNA secondary structure prediction. (**A**) Ternary plots highlighting the mismatch signature found in DMS-modified yeast rRNAs, from MaP-Seq experiments using TGIRT. For each reference nucleotide, the relative proportion of the other three nucleotides at mismatched positions is shown. Mismatches are colored according to their experimental pairing status (www.rna.icmb.utexas.edu) (**B**) Combination of mismatch frequency and mismatch signatures increases the separation of mismatches into “paired” and “unpaired” clusters, compared to using the mismatch frequency or signature alone (**C**) Accuracy of pairing status predictions of yeast rRNAs using RT drop-off (DMS-Seq), mutational profiling (MaP) or a combination of both methodologies (RT-stop + MaP). Error bars represent standard deviations. Significance: * p < 0.05; ** p < 0.01; *** p < 0.001; n.s. non-significant (**D**) ROC curves of the pairing status predictions of yeast rRNAs using RT drop-off (DMS-Seq), mutational profiling (MaP) or a combination of both methodologies (RT-stop + MaP) (**E and F**) ARC plots depicting the predicted RNA secondary structure of yeast 25s rRNA, covering the regions 2810-2952 (E) and 2977-3134 (F). RNA structure has been predicted using no experimental data (*in silico*), constraints from the RT-stop or mismatch signals, or constraints from both methodologies (see Methods). For comparison, the reference secondary structure is shown in the bottom half of each panel. Pairing interactions have been colored as follows: same pairing observed in the experimental and predicted structure (true positives, TP) in green, incorrectly predicted pairings (false positives, FP) in red, and not predicted pairings (false negatives, FN) in black. Sensitivity is defined as TP/(TP+FN); positive predictive value (PPV) is defined as TP/(TP+FP).

To test whether the inclusion of mutational signature information might improve the accuracy of prediction of DMS-probed positions identified using MaP, we trained a Support Vector Machine (SVM), and compared the accuracy of mismatches predicted using either only mismatch frequencies, or using a combination of mismatch frequencies and mutational signatures (i.e., the relative misincorporation of A, C, G, and T at each mismatch position), and find that the inclusion of mutational signature information significantly improves the accuracy of predictions of C mismatches (*p*=0.041, Student *t*-test), but not of A mismatches (*p*=0.79, Student *t*-test) (**Figure S4**).

### Combination of MaP and RT-stop produces more accurate RNA secondary structure predictions

Here we have shown that RT-stop and MaP methodologies capture non-overlapping information (Figures 1D-E and 2A-E). We therefore hypothesized that the combination of signals from both methodologies might increase the accuracy of RNA secondary structure predictions, compared to using only one of them. Here, we again used the previously published DMS-probed *S. cerevisiae* datasets ^15,25^, as these datasets include yeast rRNAs for which the RNA structure has been experimentally determined ^37^.

We first analyzed the performance of each methodology separately. As expected, we found that RT drop-off predicts RNA structures more accurately in DMS-Seq datasets compared to mutational profiling, whereas mutational profiling performs better in predicting RNA structures in DMS-MaPSeq datasets (**Figure S5**). It is important to note that the two aforementioned DMS-probed *S. cerevisiae* datasets exhibit important differences in their library preparation. DMS-Seq datasets include a second size-selection step to enrich for RT-truncated reads ^15^, and therefore should favor the detection of DMS-probed positions using RT-stop. In contrast, DMS-MaPSeq datasets do not include this second size-selection step, and, consequently, mismatch methodologies are expected to yield better results in this type of datasets. Thus, to combine signals from both methodologies, we employed RT-stop accessibilities from DMS-Seq and mutational profiles from DMS-MaPSeq.

Mutational profiling methods identify fewer DMS-probed nucleotides than RT-stop methodologies, but its positive predictive value (i.e., the proportion of true positives relative to all positives) is slightly higher (**Figure S6**). We therefore reasoned that the best possible combination of MaP and RT-stop signals might be to have MaP overrule the RT-stop accessibilities, employing the SVM pairing status predictions to select which MaP nucleotides should overrule RT-stop accessibilities. More specifically, if the SVM predicted a mismatch to be “unpaired”, the accessibility of the nucleotide was assigned to 1 (see Methods). When combining both types of information (RT-stop and MaP), we observe an average increase in accuracy of 8% with respect to RT-stop methodologies alone, and of 11% with respect to mutational profiling alone (Figure 5C). Similarly, we find that the area under the curve (AUC) of the Receiver Operating Characteristic (ROC) curves is increased when combining both methodologies (Figure 5D) (AUC values: MaP = 0.669 (rep1), 0.725 (rep2); RT-stop = 0.749 (rep1), 0.729 (rep2); combined = 0.837 (rep1), 0.834 (rep2)).

We then examined whether the increase in accuracy of predicted pairing statuses led to better RNA secondary structure predictions. To test this, we predicted the folding of 58 different domains found within the yeast 18s and 25s rRNAs using the *Fold* software ^38^ implementing the DMS signal as folding constraints (see Methods). We find that the combination of RT-stop and MaP signals yields to more accurate secondary structure prediction, mainly due to increased positive predictive values (PPV) (Figures 5E-F and **S7**). Overall, our results suggest that best RNA structure predictions are achieved when using a combination of methodologies.

## DISCUSSION

Next-generation sequencing (NGS) has revolutionized the field of molecular biology, opening new avenues to explore the genome, epigenome and transcriptome. In the last few years, genome-wide techniques to explore additional layers of regulation, such as RNA structure or the epitranscriptome, have become available. Due to their relatively recent appearance, we are still facing the challenges of determining how to best analyze these data, as well as how to properly interpret the results.

Current RNA structure studies are mainly limited to probing single-stranded (ss) regions. Upon DMS treatment, single-stranded RNA undergoes methylation in three of its four nucleotides, giving rise tom^1^A, m^3^C and m^7^G, respectively. Mutational profiling and reverse truncation signals are two methodologies used to identify DMS-modified positions. In addition to the methylations that occur upon DMS treatment, around 20 different naturally occurring RNA modifications can also affect reverse transcription^39,40^, causing either reverse transcription truncation and/or increased mismatch rates at the modified position ^29,30,35,41^. Thus, it is essential to maximize the analysis of such datasets and capture the whole spectrum of the signal.

Here we show that mutational profiling and reverse truncation both identify DMS-modified positions, however, each method captures only part of the structural information. Although there is a significant overlap of identified positions between the two methodologies (Figures 1D, 1E, 2A-C and **S1**), the correlation between positions identified using mismatch frequencies and RT drop-offs (accessibilities) is poor (Figures 2C and D). Indeed, each methodology is biased towards identifying different subsets of DMS-modified positions, being m^3^C preferentially identified by MaP and m^1^A preferentially identified by RT drop-off, in the case of using SS3 (Figure 2E). Therefore, contrary to current practice, we suggest that the optimal identification of RNA structure is generated by the union of mismatch and RT drop-off signals (Figures 5C-F).

We also show that multiple positions identified in DMS-Seq datasets as mismatches are likewise found in corresponding RNA-seq data (Figure 3C) and thus unrelated to DMS probing. Coupling RNA-Seq with DMS-Seq allows for filtering out these positions, increasing the signal-to-noise ratio. Compared to other genome-wide RNA structure probing reagents such as SHAPE, DMS is especially valuable to optimize the filtering steps as it only modifies A and C bases, allowing the assessment of the signal-to-noise ratio based on the number of predicted DMS-modified positions that fall at G and T bases. Thus, we were able to develop a robust filtering strategy to reduce the noise-to-signal ratio, increasing the accuracy of the signal (**Figure S3**). In addition, we also show that this signal can be further improved if the mutational signature is taken into consideration (**Figure S4**)

Importantly, our findings are not only applicable to RNA structure, but also to the field of RNA modifications. RNA modifications are known to alter the structure, function and activity of RNA molecules ^42–48^. Recent papers have analyzed multiple RNA-Seq datasets looking for mismatched nucleotide signatures, finding that many RNA sequences contain modified nucleosides, ^29,30,49^, and in the last year, genome-wide maps identifying thousands of m^1^A modifications have been made available ^50^. However, previous studies have mainly used mutational profiling to identify RNA modifications from these datasets ^29^. We suggest a combination of RT truncation and mutational profiling methodologies may facilitate a more complete identification of genome-wide RNA modifications.

Furthermore, our analysis shows that the choice of the RT enzyme not only affects the mismatch/RT drop-off ratio, but also the relative proportion of misincoporated nucleotides (Figure 4A-C), dramatically affecting the mismatch signatures (Figure 4D-F). Consequently, mismatch signatures obtained with a given RT enzyme (e.g., SS3) cannot be used to predict DMS-modified positions in datasets from samples reverse-transcribed with a different RT enzyme (e.g., TGIRT). This is also true for analyses aiming to detect naturally occurring RNA modifications based on mismatch signature ^29,30,49^. Whether additional variables like RT temperature or salt concentration may affect the mismatch signature of RNA modifications is still an open question.

Overall, we show the DMS signal cannot be entirely captured either qualitatively or quantitatively using only mutational profiling or RT drop-off methodologies (Figures 2B-D). Indeed, we find that mutational profiling preferentially identifies DMS-modified C bases (m^3^C) whereas RT drop-off preferentially identifies DMS-modified A bases (m^1^A) (Figures 2C and 2E). Therefore, the relative proportion of mismatch/RT drop-off is dependent on the identity of the modified base. While more processive RT enzymes, such as TGIRT, increase mismatch frequency over drop-off, we show that this increased processivity is not equal across all RNA modifications. Finally, we show that a combination of signals improves the accuracy of RNA secondary structure predictions (Figure 5). Consequently, we propose that both MaP and RT drop-off signals should be employed to obtain the most accurate predictions from genome-wide RNA structure probing datasets, regardless of the reverse transcriptase employed during the library preparation.

## MATERIALS AND METHODS

### Zebrafish maintenance

Wild-type zebrafish embryos were obtained through natural mating of TU-AB strain of mixed ages (5-18 months). Mating pairs were randomly chosen from a pool of 70 males and 70 females allocated for each day of the month. Fish lines were maintained following the International Association for Assessment and Accreditation of Laboratory Animal Care research guidelines, and approved by the Yale University Institutional Animal Care and Use Committee (IACUC).

### DMS-Seq, RNA-Seq and Structure-Seq datasets

DMS-Seq datasets of *in vitro* DMS treated known RNA structure spike-ins (*Tetrahymena* ribozyme and DsRed mRNA containing a tRNA-spinach cassette in its 3’UTR) and *in vivo* DMS treated 64c stage zebrafish embryos (samples: SRS2404542 and SRS2404544) were taken from SRP114782. RNA-Seq datasets from 64c stage zebrafish embryos (samples: SRS2404514 and SRS2404517) were also taken from SRP114782. The manuscript associated with SRP114782 is currently under review and all datasets will be made publicly available immediately after acceptance. In the meantime, all datasets related to SRP114782 are available upon request. Structure-Seq datasets of 64-cell stage zebrafish embryos using either SSIII or TGIRT reverse transcription enzymes are accessible at SRP115809. See Table S1 for more details on all datasets. DMS treated yeast rRNA data was taken from previously published datasets GSE45803 and GSE84537.

### Structure-Seq experiments

For *in vivo* modification of zebrafish embryo transcriptome, 150 64-cell embryos were transferred to 5 mL eppendorf tubes containing 400 µL of system water from the fish facility. 100% DMS (Sigma-Aldrich) was diluted in 100% ethanol to obtain a 20% DMS stock solution. The DMS stock solution was then diluted to 6% DMS in 600 mM Tris-HCl pH 7.4 (AmericanBio) in system water from the fish facility. This DMS/Tris-HCl solution was immediately mixed vigorously and 200 µL was added to each embryo containing tube to reach a final concentration of 2% DMS and 200 mM Tris HCl pH 7.4. Embryos were incubated at room temperature for 10 min with occasional gentle mixing. The DMS solution was then quickly removed from the tubes and the embryos were flash frozen in liquid nitrogen. Frozen embryos were thawed and actively lysed with 800 µL of TRIzol (Life Technologies) supplemented with 0.7 M ß-mercaptoethanol (Sigma-Aldrich) to quench any remaining trace of DMS. After 2 min incubation, TRIzol was added to reach a final volume of 4 mL and total RNA extracted following the manufacturer’s protocol. Poly(A)+ transcripts were purified using oligo d(T)_25_ magnetic beads (New England BioLabs) following the manufacturer’s protocol and eluted in 35 µL of water. DMS treatments were performed in duplicate from different clutches and days. For each replicate, an untreated control was performed following the same steps, omitting the DMS.

Structure-Seq libraries were prepared as in Ding *et al.* ^13^ with few changes. Briefly, DMS treated or untreated poly(A)+ RNA duplicates were pooled together and subjected to reverse transcription using a partially degenerated primer fused with part of an Illumina TruSeq adapter (5’-AGACGTGTGCTCTTCCGATCTNNNNNN-3’) and either the SuperScript III First Strand Kit (Invitrogen) or the TGIRT^TM^-III (InGex) reverse transcriptase following manufacturer’ protocols. Each type of reverse transcription reaction was performed using the same pool of DMS-modified RNAs allowing a direct comparison of the two enzymes. For reverse transcription reactions with the SuperScript III enzyme, samples were heated at 25°C for 10 min, 42°C for 30 min, 50°C for 10 min, 55°C for 20 min, and 75°C for 15 min to deactivate the enzyme. For reverse transcription reactions with the TGIRT^TM^-III enzyme, samples were heated at 25°C for 10 min, 42°C for 10 min, 50°C for 10 min, 55°C for 10 min, 60°C for 30 min, 65°C for 20 min, and 75°C for 15 min to deactivate the enzyme. All samples were treated with RNAse H at 37°C for 20 min. cDNAs were purified using 36 µL of Agencourt AMPure XP beads (Beckman Coulter) following manufacturer’ protocol and resuspended in 10 µL of water. ssDNA linker (/5Phos/NNNNNGATCGTCGGACTGTAGAACTCTGAAC/3InvdT/) was ligated at cDNA 3’-ends using the CircLigase ssDNA ligase (Epicentre) with slightly modifications to the manufacturer’s protocol, i.e., reagents were added to 3 µL of cDNAs to reach the following final concentrations: 50 units of CircLigase, 0.05 mM ATP, 2.5 mM MnCl_2_, 10% PEG 6000, 1 M betaine, and 5 µM ssDNA linker in a volume of 10 µL. Ligation reactions were incubated at 60°C for 2h, 68°C for 1h, and 80°C for 10 min to deactivate the ligase. 10 µL of water was added to each reaction. The resulting 20 µL ligation products were purified using 36 µL of Agencourt AMPure XP beads and dissolved in 16 µL of water. PCR amplification was performed on the ligated cDNA using Illumina primers (Small RNA PCR Primer 2 5’-AATGATACGGCGACCACCGACAGGTTCAGAGTTCTACAGTCCGA-3’ and PCR primer index 5’-CAAGCAGAAGACGGCATACGAGATbarcodeGTGACTGGAGTTCAGACGTGTGCTCTTCCGATCT-3’, where barcode is the 6-nucleotide index). PCR products were purified and concentrated using the MinElute PCR Purification Kit (QIAGEN) and eluted in 10 µL of water. Eluted PCR products were separated in a 2% agarose gel and products of 200-1,000 nucleotide were extracted and purified using the MinElute Gel Extraction Kit (QIAGEN). Libraries were sequenced on Illumina HiSeq 2000/2500 machines producing single-end 76 nucleotide reads. Sequencing samples are summarized in Table S1.

### Read filtering and mapping

DMS-seq raw reads contained the following features: NNNN-insert-NN-barcode(4-mer)-adapter where the 6N (NNNN+NN) sequence composes the Unique Molecular Identifier (UMI), “barcode” is the sample 4-mer in-house barcode and adapter is the 3’-illumina adapter. The UMI was used to discard PCR duplicates and count single ligation event. The barcode was used to mark individual replicates following the 3’-adapter ligation step. Base calling was performed using CASAVA-1.8.2. The Illumina TruSeq index adapter sequence was then trimmed by aligning its sequence, requiring 100% match of the first five base pairs and a minimum global alignment score of 60 (Matches: 5, Mismatches: −4, Gap opening: −7, Gap extension: −7, Cost-free ends gaps). Trimmed reads were demultiplexed based on the sample’s in-house barcode, the UMI was clipped from the 5’- and 3’-end and kept within the read name, for marking PCR duplicates. Structure-Seq reads contained the following features: NNNNN-insert, where 5N correspond to the UMI, and were processed as for the DMS-Seq reads omitting the Illumina index adapter trimming and clipping only the 5’-end UMI. DMS-Seq and Structure-Seq reads were then depleted of rRNA, tRNA, snRNA, snoRNA and miscRNA, using Ensembl78 annotations, as well as from RepeatMasker annotations, using strand-specific alignment with Bowtie2 v2.2.4 ^51^. The remaining reads were aligned to the zebrafish Zv9 genome assembly using STAR version 2.4.2a ^52^ with the following non-default parameters: *--alignEndsType EndToEnd --outFilterMultimapNmax 100 --seedSearchStartLmax 15 --sfbdScore 10 -- outSAMattributes All*. Genomic sequence indices for STAR were built including exon-junction coordinates from Ensembl 78. Only reads of unique UMI were kept at each genomic coordinate for DMS-seq and ribosome profiling experiments. Raw reads from RNA-seq experiments were processed using the same pipeline, omitting the adapter trimming, barcoding demultiplexing and UMI clipping steps. The filtered reads were aligned onto Zebrafish Zv9 assembly using STAR, with the same parameters as described above. STAR genomic sequence indices were built including exon-junction coordinates from Ensembl 78.

### Analysis of accessibility (RT-stop methodology)

Per-transcript profiles were computed using uniquely mapped reads overlapping at least 10 nucleotides with the transcript’s annotation. Each read count was attributed to the nucleotide in position –1 of the read’s 5’-end within the transcript coordinate, to correct for the fact that reverse transcription stops one nucleotide prior to the DMS-modified nucleotide. To determine read distributions for each nucleotide, only transcripts with a minimum of 100 counts were considered. Accessibilities were calculated following the 2%-8% rule ^53^, i.e., by normalizing the read counts proportionally to the most reactive As and Cs within the region after the removal of outliers. More specifically, the 2% most reactive As and Cs were discarded and each position was divided by the average of the next 8% most reactive As and Cs. Accessibilities greater than 1 were set to 1, and accessibilities for G and T were set to 0.

### Analysis of mutational profiles (MaP methodology)

DMS-Seq, Structure-Seq and RNA-Seq mapped bam files were processed using HAMR^29^ to produce a bed file of mismatched positions, including metadata information regarding reference nucleotide, coverage (number of reads at each base pair), and relative nucleotide frequencies at each position (measured as total number of A, C, G, and T nucleotides, normalized by the coverage). Mismatched positions were then filtered to remove SNPs, naturally occurring RNA modifications and sequencing artifacts (see below). Due to the low mismatch frequencies that m^1^A and m^3^C modifications cause, a minimum of 50 reads/bp was required. Overall, the set of filtered mismatch positions met all the following criteria: i) found in both DMS-Seq replicates; ii) not found in the set of replicable RNA-Seq mismatches; iii) minimum coverage of 50 reads/bp; iv) minimum of 2 mismatched reads/bp; and v) maximum 0.25 mismatch frequency. For Structure-Seq datasets, the set of mismatched positions was filtered by the control (untreated) Structure-Seq mismatches, instead of the RNA-Seq mismatches.

### Incorporation of mutational signatures to increase the accuracy of mismatched nucleotides (MaP methodology)

Mutational signatures, defined as the proportion of misincorporation of each nucleotide (i.e., number of As, Cs, Gs, Ts, at each mismatched position), were used to further refine the prediction of whether a position with increased mismatch rate was a true or false positive. Yeast rRNA mismatches for which the experimental structure is known were used to train a Support Vector Machine (SVM). The data was subdivided into training (25%) and testing (75%), using the latter to compute the accuracy of the SVM predictions (**Figure S4C**). Five-fold cross-validation was used to determine the evaluate the SVM prediction model. ROC curves were computed using the *ROCR* package in R. Ternary plots were generated using the *ggtern* package in R. SVM training was performed using the *e1071* package in R. The coefficient of overlap of distributions was computed using the *overlap* and *hypervolume* packages in R.

### Accuracy of pairing status predictions of yeast rRNAs

To determine the accuracy of each methodology (RT-stop, MaP, and RT-stop + MaP), the accessibilities, mismatch frequencies, or combination of both signals of yeast rRNAs were used to train a Support Vector Machine (SVM), respectively. For this aim, the data was subdivided into training (25%) and testing (75%), using the latter to compute the accuracy of the SVM predictions, for each methodology (Figure 5C).

### Combination of RT-stop and MaP signals

The MaP methodology identifies fewer DMS-probed nucleotides than the RT-stop methodology, but these nucleotides tend to be enriched in true positives (**Figure S6**). Therefore, we combined the signals by forcing a position to be unpaired if MaP identified it as unpaired. More specifically, if the SVM (which employs a combination of mismatch frequency and mutational signatures to predict the outcome) predicted a mismatch to be “unpaired”, the accessibility of the nucleotide was assigned to 1. RT-stop information was computed from previously published *in vitro* probed yeast DMS-Seq datasets (reverse-transcribed using SS3) and yeast MaPSeq datasets (reverse transcribed using TGIRT), corresponding to GSE45803 and GSE84537, respectively. Mutational profiles generated using TGIRT were chosen for combining signals, as we found that these profiles were much more accurate than those generated with SS2 (AUC=0.71 using TGIRT; AUC=0.59 using SS2) (**Figure S5**), in agreement with previous works ^25^. Experimentally yeast rRNA structures were obtained from www.rna.icmb.utexas.edu. The predicted secondary structure of 58 different domains found in the 18s and 25s rRNAs was computed using the *Fold* executable^38^ of the RNA structure package (version 5.6)^54^ with the following parameters: -*mfe* and –*t 303.15*. The DMS signal was used as soft constraint with the –*dms* option. Each predicted structure was compared to the reference structure using the scorer executable from the same package to obtain the sensitivity and positive predictive value (PPV). Arc plots representing RNA secondary structure were drawn using the R-chie software^55^ with default parameters. It is important to note that in order to combine RT-stop and drop-off, maximal performance is achieved when using two different libraries: RT-stop from DMS-Seq and mutational profiles from DMS-MaPSeq (as shown, for example, in Figure 5D). If a single library type is employed (i.e. either DMS-Seq or DMS-MaPSeq, but not both), only small improvements will be obtained, because each library type favors the performance of one of the methods (**Figure S5**).

### Code availability

In-house scripts used in this work are available at https://github.com/enovoa/dmsseq_dmsmapseq

## AUTHOR CONTRIBUTIONS

EMN and JDB designed the project. EMN performed the bioinformatic analyses, with the contribution of JDB. JDB prepared the DMS-Seq and Structure-Seq libraries. EMN and JDB interpreted the results and built the figures. AJG, JSM and MK supervised the project. EMN and JDB wrote the paper, with the contribution of all authors.

## ACKNOWLEDGEMENTS

We thank all members of the Giraldez, Mattick, and Kellis labs for their valuable comments and suggestions. This research was supported by the Human Frontier Science Program (LT000307/2013-L to EMN), the Australian Research Council (DE170100506 to EMN), the Fonds de Recherche du Québec - Santé (postdoctoral fellowship to JDB), the National Health and Medical Research Council (Project Grant APP1070631 to JSM) and the National Institute of Health (grants R01 HD074078, GM103789, GM102251, GM101108 and GM081602), Pew Scholars Program in the Biomedical Sciences, March of Dimes 1-FY12-230, the Yale Scholars Program and Whitman fellowship funds provided by E. E. Just, Lucy B. Lemann, Evelyn and Melvin Spiegel, The H. Keffer Hartline and Edward F. MacNichol, Jr. of the Marine Biological Laboratory in Woods Hole, MA to AJG.

