## Supplementary Figures for "Best practices for genome-wide RNA structure analysis: combination of mutational profiles and drop-off information"

**Figure S1. RT-stop and MaP DMS signals are in agreement with known RNA structures.** (**A and B**) Comparison of the overlay of accessibilities calculated from RT-stop (left) and mismatch frequencies (right) onto two different regions of the experimentally determined RNA secondary structure of the *Tetrahymena* ribozyme. Each nucleotide has been colored based on its normalized accessibility or mismatch frequency, using the 2%-8% normalization. Arrows point to examples where the nucleotide is: *i*) preferentially predicted by the RT-stop method (green), *ii*) preferentially predicted by mutational profiling (red), or *iii*) equally predicted by both methods (orange) (**C and D**) Box plots of the agreement between accessibilities calculated from RT-stop or mismatch signals, and A/C pairing status (ss, single-stranded; ds, double-stranded) for the *Tetrahymena* ribozyme (C) and the tRNA-Spinach cassette (D).

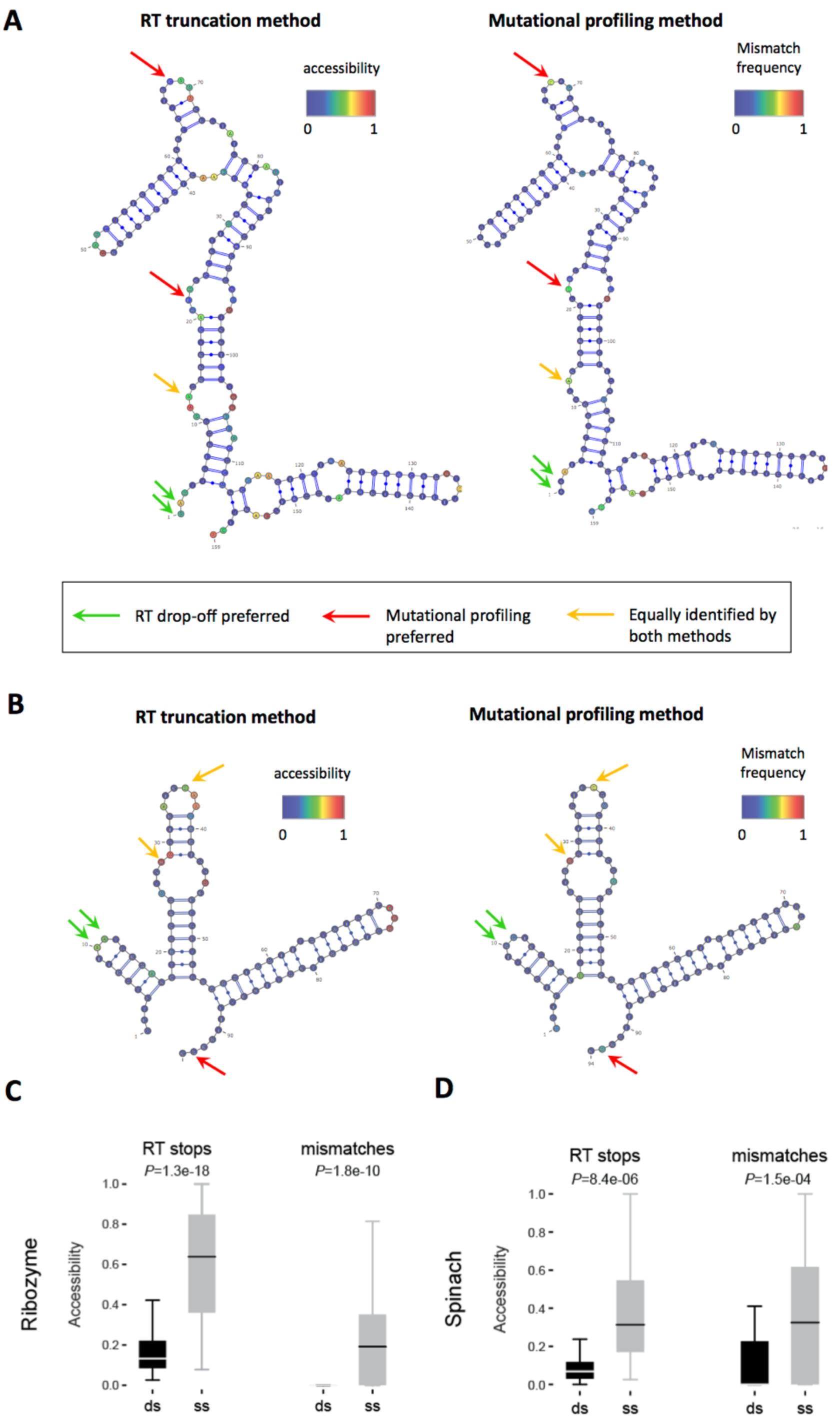

**Figure S2. Analysis of the mutational signature of the two spike-ins**. (**A and B**) Mismatch signature ternary plots for the two spike-ins, the *Tetrahymena* ribozyme and Spinach tRNA cassette, prior to (A) and after filtering (B)**.** For filtering details, see Methods. (**C and D**) Relative proportion of misincorporated nucleotides at mismatched positions, when the reference nucleotide is A (top) or C (bottom) in the spike-ins (C) and in the transcriptome-wide DMS-probed 64-cell stage zebrafish embryos (D).

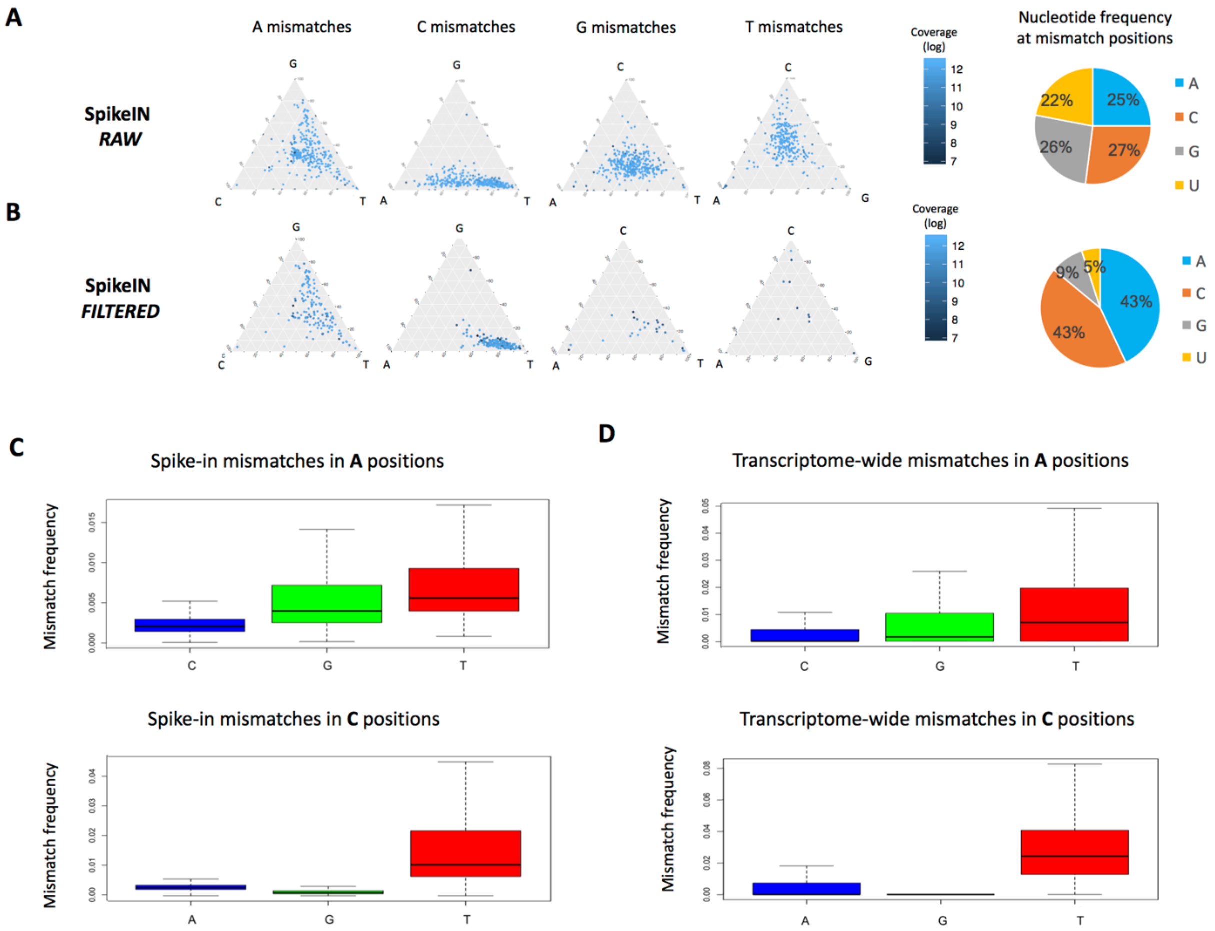

**Figure S3. Filtering methodology to capture high confidence DMS-MaPSeq signal in a vertebrate transcriptome.** Mismatch signature plots of 64-cell zebrafish embryos transcriptome-wide DMS-Seq samples, for each reference nucleotide, after each step of filtering. The relative proportion of the other three nucleotides at mismatched positions is shown as a ternary plot. Ternary plots have been computed at each stage of filtering. The proportion of reference nucleotides with mismatches after each filtering step is depicted on the right side of the figure.

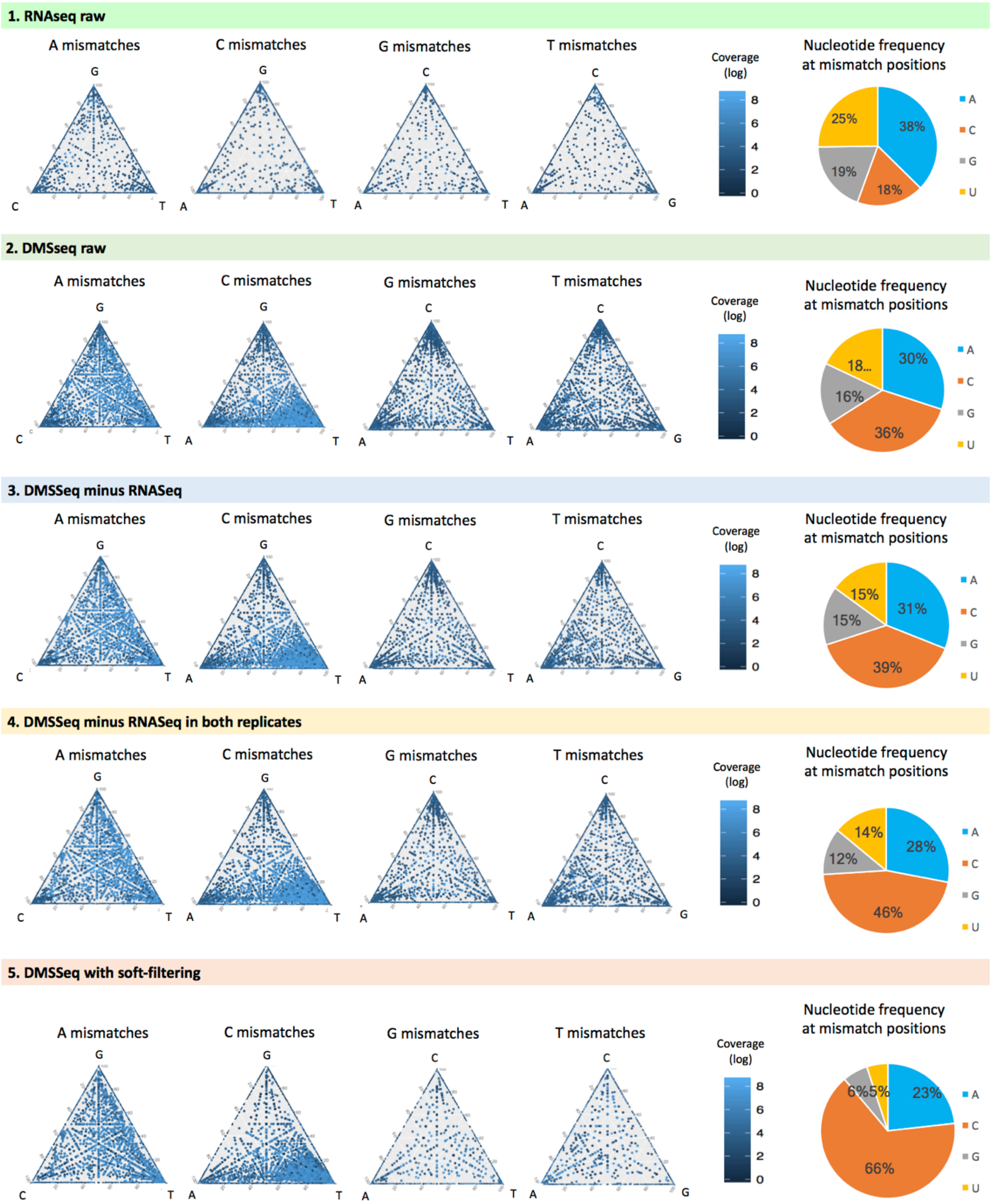

**Figure S4. Combination of mismatch signatures and mismatch frequencies improves the accuracy of RNA pairing status predictions (A and B)** Correlation between mismatch signatures (T/G ratios for A mismatches; T/A ratios for C mismatches) and mismatch frequencies in yeast rRNAs, using MaPSeq datasets reverse transcribed using TGIRT. Each position is colored according to either its experimentally determined pairing status (www.rna.icmb.utexas.edu) (A) or predicted pairing status after SVM training (B). (**C**) Accuracy between the predicted pairing status and the experimentally determined pairing status. The predicted pairing status was determined using either mismatch frequencies (orange), or a combination of mismatch frequencies and mutational signatures (gray). Significance: * $p < 0.05$; n.s. non-significant.

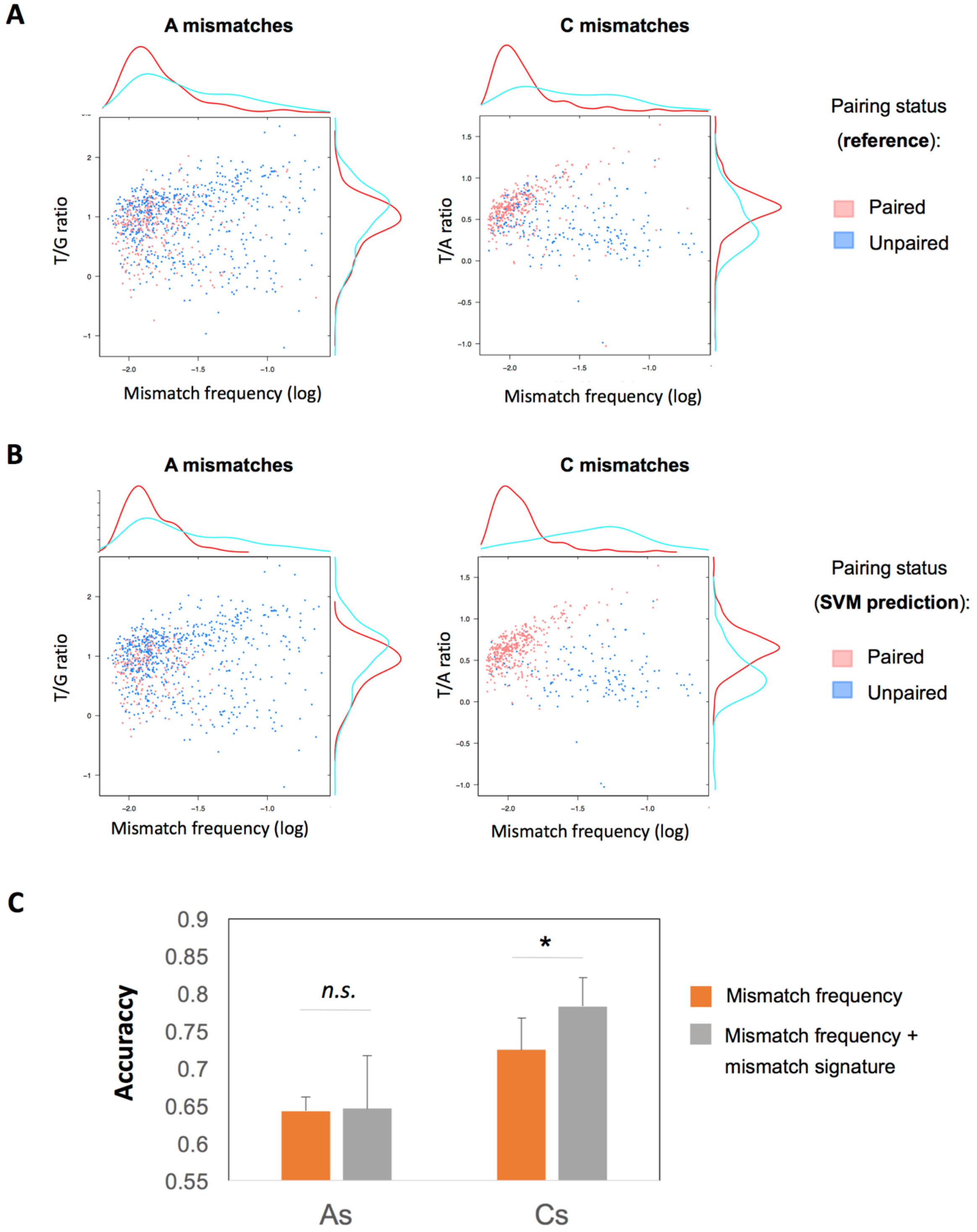

**Figure S5. Comparison of the performance of MaP and RT-stop methodologies, in DMS-seq and DMS-MaPSeq datasets.** ROC curves depicting the accuracy of RNA structure pairing status predictions of yeast rRNAs, using either DMS-MaPSeq datasets (top panels) or DMS-Seq datasets (bottom panels). For each dataset type, pairing statuses were computed using mutational profiling (red) and RT-drop off (blue). Each biological replicate is plotted independently.

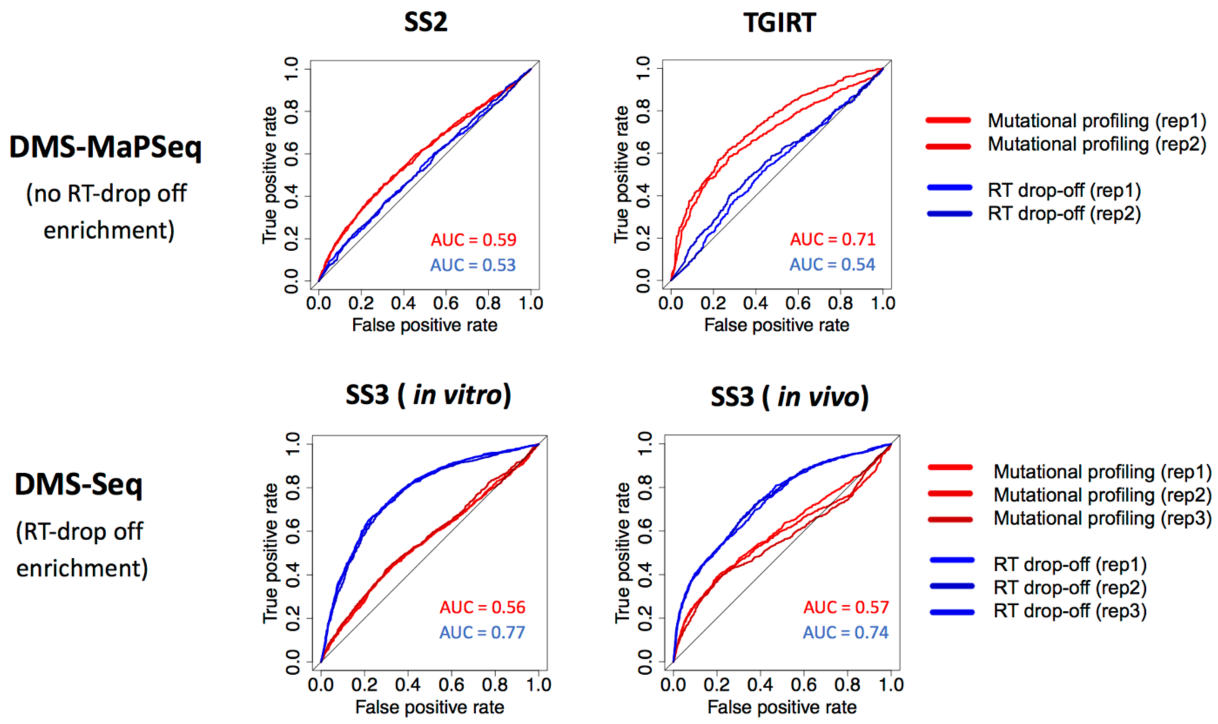

**Figure S6. Distribution of accessibility and mismatch frequency values in yeast rRNAs.** Positions have been binned according to their pairing status. To subdivide all positions into correctly and incorrectly predicted for each pairing status (i.e., TP, FP, TN, FN). Threshold is depicted with a red horizontal line. Positive predictive values (PPV), which represent the proportion of true positives (TP) relative to the total of positives (TP+FP), are shown for each of the two methods, and for each replicate. The DMS-MaPSeq dataset reverse transcribed using TGIRT was used to obtain the mismatch frequencies depicted in the MaP method.

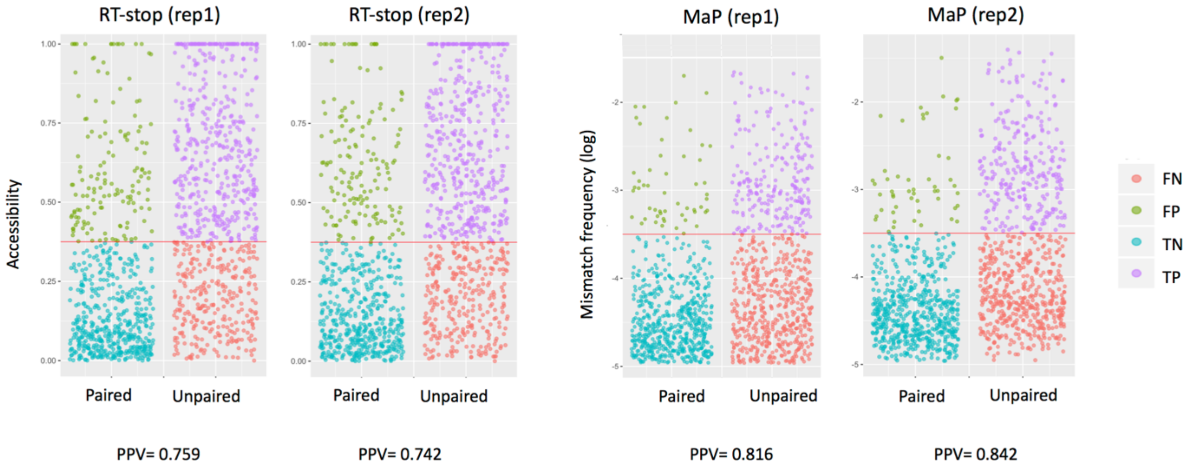

**Figure S7. Sensitivity and PPV boxplots of the RNA secondary structure predictions for different domains from the 18s and 25s rRNAs**. RNA secondary structure has been predicted using the *Fold* software, without constraint (in silico, gray) or with DMS-based constraints (RT-stop (red), MaP (blue) and RT-stop+MaP (green)). From the 58 domains analyzed, only those with *in silico* sensitivity and PPV below 0.5 were selected for further analysis (n=20 and n=25 domains, respectively), to allow for improvement. *P*-values were computed using Wilcoxon signed ranked test (n.s. = non-significant).

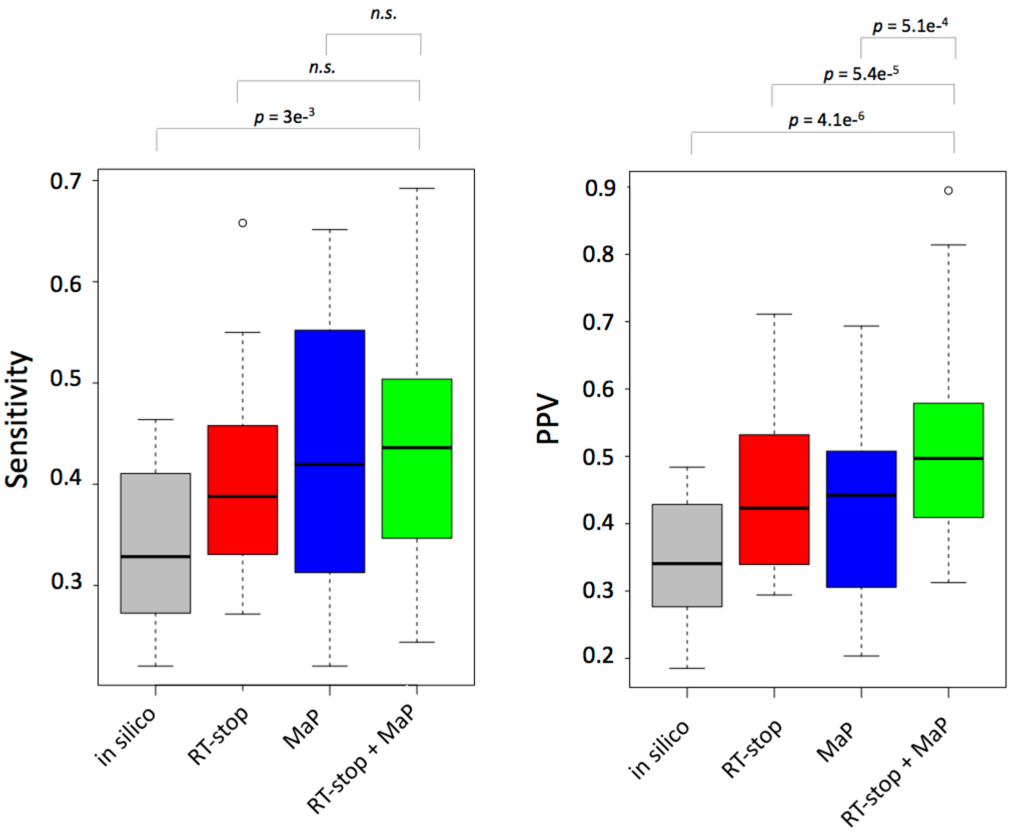
